# Sonic restoration: Acoustic stimulation enhances soil fungal biomass and activity of plant growth-promoting fungi

**DOI:** 10.1101/2024.01.11.575298

**Authors:** Jake M. Robinson, Christian Cando-Dumancela, Martin F. Breed

**Affiliations:** College of Science and Engineering, Flinders University, Bedford Park, SA 5042, Australia; The Aerobiome Innovation and Research Hub, College of Science and Engineering, Flinders University, Bedford Park, SA 5042, Australia

**Keywords:** ecoacoustics, acoustic restoration, fungi, soil biodiversity, sonic restoration, soil health

## Abstract

Ecosystem restoration interventions often utilise visible elements to restore an ecosystem (e.g., replanting native plant communities and reintroducing lost species). However, using acoustic stimulation to restore ecosystems has received little attention. Our study aimed to (a) investigate the potential effects of acoustic stimulation on fungal biomass and organic matter decomposition, which are both crucial components of ecosystem functioning and (b) assess the effect of acoustic stimulation on the growth rate and sporulation of the plant growth-promoting fungus *Trichoderma harzianum*. We played 70 dB and 90 dB soundscape treatments (@ 8 kHz) to green and rooibos teabags in compost in experimental mesocosms for 8 hours per day for 14 days to test whether acoustic stimulation affected fungal biomass and organic matter decomposition (a control mesocosm received only ambient sound stimulation <30 dB). We played a monotone soundscape (80 dB @ 8 kHz) over five days to *Trichoderma harzianum* to assess whether this stimulation affected the growth rate and sporulation of this fungus (control samples received only ambient sound stimulation <30 dB). We show that the acoustic stimulation treatments resulted in increased fungal biomass, greater decomposition, and enhanced *T. harzianum* conidia (spore) activity compared to controls. These results indicate that acoustic stimulation influences soil fungal growth and potentially facilitates their functioning. A piezoelectric effect and/or fungal mechanoreceptor stimulation are possible mechanisms. Our study highlights the potential of acoustic stimulation to alter important functional soil components, which could, with further development, be harnessed to aid ecosystem restoration.

## Introduction

Ecosystem restoration is imperative in the face of escalating ecosystem degradation and global biodiversity loss (Tedesco et al. 2023). Efforts to restore ecosystems often focus on physical and visible interventions, such as revegetation (LázaroGonzález et al. 2023) and species reintroductions (Hugron et al. 2020). While these approaches are crucial for ecosystem recovery, there remains a notable gap in our understanding of how audible domains could aid ecosystem recovery, particularly below-ground. This subterranean focus is particularly important as 59% of the world’s biodiversity lives in soil (Anthony et al. 2023). Moreover, soil fauna such as earthworms, are major contributors to ecosystem functioning and food production (Fonte et al. 2023). The potential importance of audible domains in restoration invites questions about whether acoustic stimulation (the application of sound to a particular ecological receptor) could directly promote the restoration of soil ecosystems.

Ecological acoustic surveys or ‘ecoacoustics’ have proven successful at monitoring soil biodiversity (Maeder et al. 2022), which is a vital but challenging-to-monitor ecosystem component. Recently, Robinson et al. (2023) demonstrated that it is possible to record soniferous species below-ground using piezoelectric microphones and audio recording devices in a restoration context. The authors built acoustic indices of audible soil diversity, complexity and normalised differential signals that reflected the recovery of soil biodiversity in a temperate forest context. Moreover, Görres and Chesmore (2019) used similar acoustic technology to detect scarab beetle larvae stridulation in a soil pest monitoring setting.

However, the role of acoustic stimulation in fostering ecosystem recovery remains underexplored. The emerging field of ‘acoustic restoration’ aims to broadcast soundscapes in disturbed areas to facilitate the recolonisation of animals, microorganisms, and biogenic compounds (Znidersic et al. 2022). For instance, McAfee et al. (2022) enriched marine soundscapes to enhance recruitment and habitat building on oyster reefs. They deployed low-cost marine speakers at four sites and compared oyster recruitment rates. The authors found that soundscape playback significantly increased oyster recruitment at 8 of the 10 study sites.

Sound, as a fundamental aspect of the environment, holds immense potential to influence ecological processes and shape ecosystem dynamics. Similarly, anthropogenic sounds can alter ecosystem dynamics (Kunc and Schmidt, 2019). However, the impact of sound on the restoration of degraded ecosystems, particularly soil environments, has received little attention. According to a recent review (Robinson et al. 2021), studies have shown that acoustic stimulation using monotonous anthropogenic sound can change the community composition, growth rate and biomass of lab-grown bacteria (Gu et al. 2016), algae (Cai et al. 2016) and fungi (Hofstetter et al. 2020). However, there have been no studies on the effect of anthropogenic sound exposure on the recovery of soil environments or the activity of plant growth-promoting microbiota. This knowledge gap presents an opportunity to explore the relationship between acoustic stimulation and ecosystem restoration, particularly how it affects functional ecological components (e.g., biomass, diversity, plant growth/health-promoting microbiota).

Two essential ecosystem functions that are influenced by soil microorganisms are nutrient cycling (including decomposition and biomass) and plant-soil microbial interactions (Dantas de Paula et al. 2021). Soil microorganisms, including bacteria, viruses, fungi and others, drive these fundamental ecosystem processes (Wagg et al. 2019), yet their response to acoustic stimulation remains underexplored.

Investigating the potential effects of acoustic stimulation on soil fungal biomass, organic matter decomposition and plant growth-promoting activity (along with microbial community dynamics) could provide valuable insights that eventually aid ecosystem recovery.

We sought to take the first steps in understanding whether different soundscape parameters could affect soil fungal biomass, organic matter decomposition and plant growth-promoting fungal activity. To do this, we aimed to: (a) investigate the potential effects of acoustic stimulation on fungal biomass and organic matter decomposition (both key components of ecosystem functioning), and (b) assess the effect of acoustic stimulation on the growth rate and sporulation of the plant growth-promoting fungus *Trichoderma harzianum*. To examine the first aim, we played 70 dB and 90 dB soundscape treatments (@ 8 kHz) to green and rooibos teabags in compost in experimental mesocosms for 8 hours per day for 14 days (a control mesocosm received only ambient sound stimulation <30 dB). To explore the second aim, we played a monotone soundscape (80 dB @ 8 kHz) over five days to *Trichoderma harzianum* (control samples received only ambient sound stimulation <30 dB).

Understanding soil microorganism responses to acoustic stimulation could have far-reaching implications for ecosystem restoration strategies. While we aim to conduct comprehensive follow-up studies with refined soundscape parameters and detailed microbiomics techniques (e.g., deep sequencing soil microbiomes to determine functional responses), the objective of this study was to establish the foundations.

## Methods

### Experimental setup

The acoustic stimulation of soil was conducted in dedicated, sound-attenuated spaces in Hampshire, UK, between March 11 and 25, 2023. The spaces were 1.5 m x 1.5 m x 2.5 m. We sterilised the spaces using a 1% Virkon solution to prevent fungal contamination. Sound attenuation foam was installed on each wall of the study spaces to (a) reduce ambient noise and (b) prevent the controlled acoustic stimuli from escaping. Recording acoustic samples in ambient conditions may capture sounds from variable detection spaces. To address this and create controlled conditions, we built and installed three sound attenuation chambers (one per treatment) within these study spaces. The sound attenuation chambers (Figure S1) were made from heavy-duty 80 L plastic containers with secure lids and Advanced Acoustics (305 mm) Wedge acoustic studio foam installed on each internal wall of the container using Velcro strips.

The acoustic stimulation of the plant growth-promoting fungus *T. harzianum* was done in a lab at Flinders University, South Australia between December 15, 2023 and January 2, 2024. The same style of 80 L sound attenuation chambers were used. Both lab spaces were kept at a constant 25°C and the local environment was monitored with a ThermoPro TP50 digital indoor thermometer.

### Compost, teabags and containers

To establish a self-contained collection of organic matter and measure its decomposition rate, we applied an adapted version of the Keuskamp et al. (2013) Tea Bag Index. This is a standardised method for gathering data on decomposition rate and litter stabilisation in soil. The index has been tested for sensitivity and robustness in contrasting ecosystems, confirming its applicability to a wide range of conditions. The index involves using commercially available tetrahedron-shaped teabags with sides of 5 cm, containing approximately 3 g of tea. The teabag mesh size in our study was 0.25 mm, allowing microorganisms and mesofauna to enter the bag while excluding macrofauna. To standardise baseline weights, we used a scalpel to cut an incision (3-4 mm) in the teabag margin to release a small amount of leaves. This allowed us to have a consistent baseline weight of 2.8 g (measured with a Bonvoisin Digital Lab Scale with a 0.01 g accuracy).

We used two teabags per experimental unit, comprising 1 x rooibos (EAN 5060136750113) and 1 x green (EAN 5060136750052) teabag and placed them into the base of 30 x ZYBUX 6 cm round fibre plant pots (= total 120 teabags in 60 plant pots). We used potting compost in an 80 L container and measured its pH and moisture before use. We divided the contents into two: 40L was heat-treated in an oven at 100°C for 1 hr (Hawkes et al. 2002) to kill soil microorganisms and mesofauna, and the other 40L remained untreated. The heat-treated units allow for greater confidence in attributing potential changes in teabag mass or decomposition to the influence of soil biota. We then filled 30 of the plant pots with untreated compost (covering the teabags and filling to the top of the pots) and 30 with heat-treated compost. We placed the pots into the sound attenuation chambers (Figure 1) and applied different acoustic stimulation treatments (described in the next section) for 14 days. After the acoustic stimulation period, we immediately measured and recorded the weight of each teabag, heat-dried the teabags (to exclude moisture) at 70°C for 48-hrs and re-weighed them. We recorded soil pH at the beginning and end of the experiment using a Hanna GroLine Tester (China; IC-HI981030).

**Figure 1.**
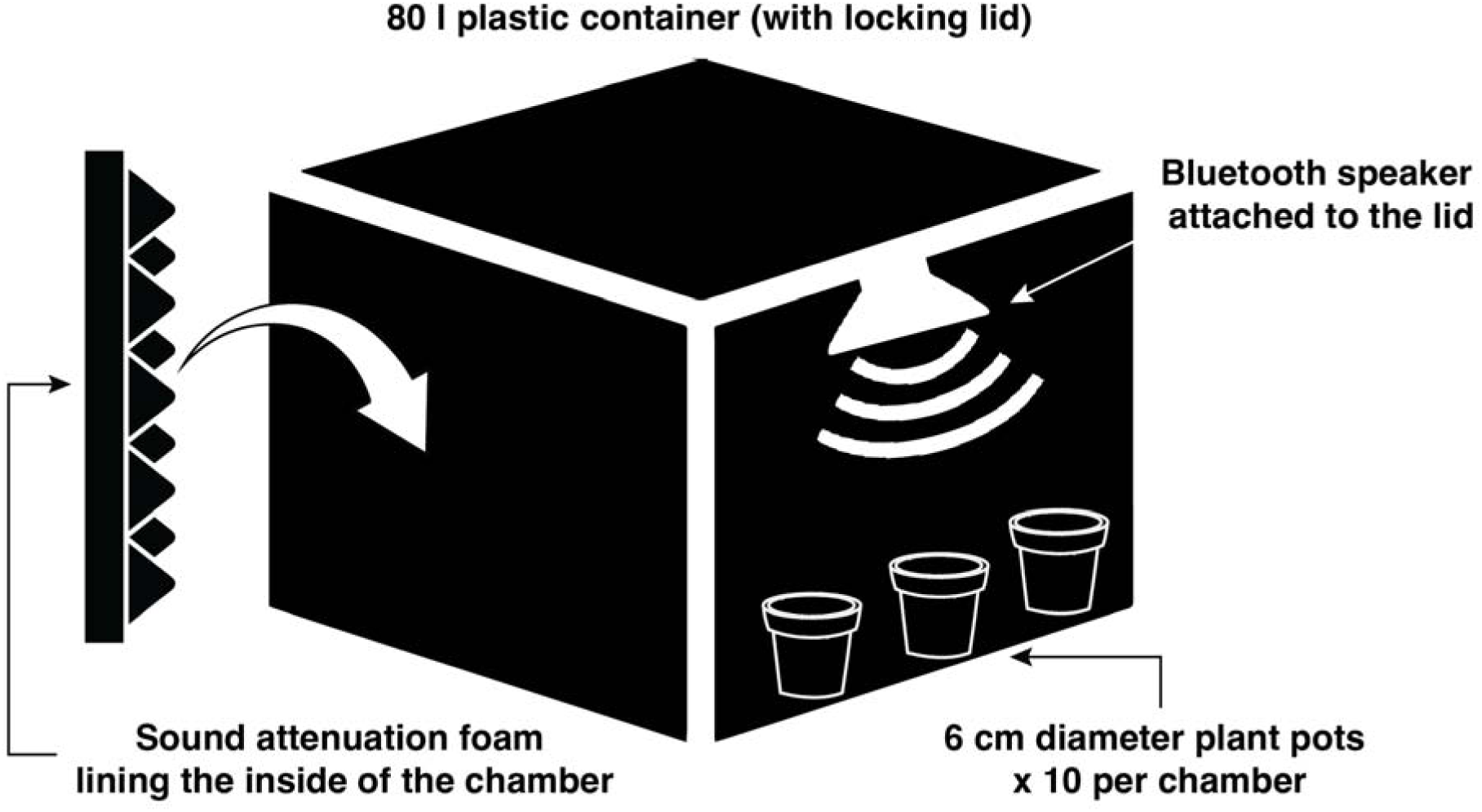
Sound attenuation chamber with pots in the base.

### Acoustic stimulation

We applied three acoustic treatments to 10 pots for the heat-treated and untreated soils in our study (Table 1). We decided upon 8 kHz as a suitable test frequency following a review (Robinson et al. 2021) that identified microbial biomass and growth rate can be influenced by this frequency. An amplitude of 80 dB is known to influence *Escherichia coli* bacteria (Gu et al. 2016), and *Chlorella* algae biomass (Jiang et al. 2012), and 90 dB influences *Picochlorum oklahomensis* (Cai et al. 2016). We used this as a guide and applied 70 dB (to capture potential lower amplitude responses) and 90 dB amplitude levels. Both amplitude levels were played at 8 kHz.

To facilitate acoustic stimulation, we downloaded (from YouTube) an 8-hour video playing a monotonous 8 kHz sound (= Tinnitus Flosser Masker at 8 kHz by Dalesnale). We tested the frequency using a Wildlife Acoustics Echometer Touch Pro bat detector (USA), designed to capture high-frequency acoustic signals. We installed an Anker Soundcore Bluetooth speaker (USA; A3102) on the inside of the sound attenuation chamber lid (using 3 x Velcro strips) with the speaker facing downwards. One Anker Soundcore speaker per sound attenuation chamber was used (= 3 in total). We connected the Bluetooth speakers to 3 x Lenovo Tabs (China; M8) to play the sound continuously for 8 hrs each day for 14 days, starting at approximately 08:00. To determine the amplitude level in the sound attenuation chambers, we used a Uni-T Professional Meter (China; TUT352) with an amplitude detection range of 30 dB to 130 dB and adjusted the tablet sound accordingly.

### *T. harzianum* culture assay

We selected *T. harzianum* as our focal plant growth-promoting fungus for three reasons: (a) it has several potential beneficial functions that could enhance ecosystem restoration (e.g., P solubilisation, ability to synthesise beneficial phytohormones, and ability to outcompete plant pathogens) (Li et al. 2015; Illescas et al. 2021; Swain and Mukherjee, 2020), (b) it is not an obligate symbiont, and is therefore relatively easy to culture, and (c) it produces vivid green conidia (spores) that can enhance the quantification process. We used *T. harzianum* (Isolate Td22;

Organic Crop Protectants) and created a modified potato dextrose agar culture medium with 125 g potato, 15 g dextrose, 10 g baker’s yeast extract and 850 ml of distilled water (Jahan et al. 2013). The medium was created in aseptic conditions and poured/set under a laminar flow hood (Lab Systems). We combined 5 g of the *T. harzianum* per litre of distilled water to create a suspension and homogenised by shaking/swirling the flask for 30 s. We then used a sterile loop to inoculate the culture medium with *T. harzianum* in a random order, placing one small circular streak (5 mm diameter) in the centre of the Petri dish (again, under a laminar flow hood). This allowed for efficient mycelium radial growth measurements. The Petri dishes (*n* = 20 for the acoustic stimulation treatment and *n* = 20 for the control group) were then sealed with Parafilm and placed into their respective sound attenuation chambers using a digital randomiser.

### Acoustic stimulation of *T. harzianum*

We decided to average the amplitude levels from the first part of the study (i.e., 70 dB and 90 dB) to provide acoustic stimulation parameters of 80 dB @ 8 kHz. This was applied to 20 Petri dishes (in the acoustic stimulation treatment group only) using a randomised controlled trial design. We randomly selected Petri dishes to be stimulated for 30 minutes each per day, so that all 20 dishes received stimulation in a random order. This was repeated over 5 days. We used three sound attenuation chambers: one to store the acoustic-stimulation treatment group, one to store the control group (no stimulation), and one to use for the Petri dishes isolated for stimulation – these were placed directly on the Bluetooth speaker, which was on the base of the chamber facing upwards. We used the same amplitude meter and tablet for the sound source used for the first study aim.

### *T. harzianum* radial growth and conidia density quantification

We measured the radial growth of the *T. harzianum* mycelium in each Petri dish each day using a standard ruler and noted down the diameter at four points to get an average diameter in mm. We also used a novel raster analysis approach in Python (described in *Statistics and data analysis* below). To measure *T. harzianum* conidia (spores) density, we poured 10 ml of distilled water over each Petri dish after 5 days and collected the fungal biomass in 15 ml centrifuge tubes (Figure 2A-F). We then filtered out non-target fungal biomass in each sample using a sterilised sieve with a 50 μm pour size (Retsch 41105003 Testsigter) and retained the suspension containing the conidia. These were stored at 4°C. We inoculated a haemocytometer (Ozlab, Neubauer-improved, 0.1 mm depth) with 1 μl of the conidia suspension and covered the well with a cover slip (Figure 2G). We used a microscope (Wild M3 stereo) to count the cells in the four corner squares and the central square of the haemocytometer, as per standard protocols (Abdulmalik et al. 2023; Milan et al. 2024). The suspensions were diluted by 10 x to reduce the conidia density enough for quantification.

**Figure 2.**
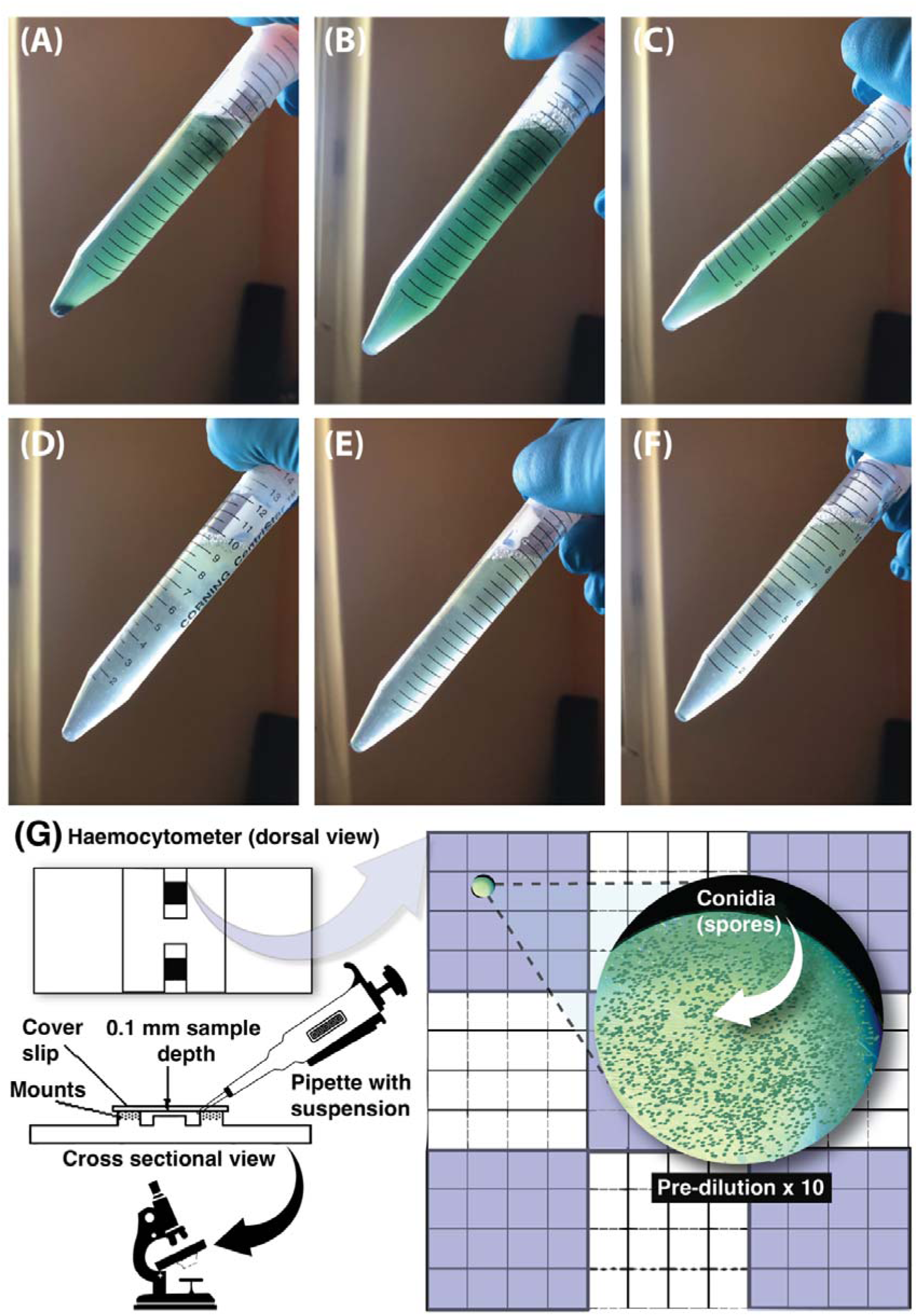
Three randomly selected conidia suspension tubes from each treatment group (A, B and C from the acoustic stimulation group, and D, E, and F from the control group) and (G) haemocytometer methods for counting *T. harzianum* conidia.

### Statistics and data analysis

Statistics were conducted in R Version 4.3.1 in R Studio 2023.06.0 “Beagle Scouts” (R Core Team, 2023) and Python (version 3.12) with supplementary software (e.g., Microsoft Excel for .csv file processing). We used ANOVA using the easyanova package in R (Arnhold 2022) to assess the effects of heat-treated soil (= treated or untreated) and acoustic treatments (= baseline, ambient, 70 dB, 90 dB) on organic matter weight, applying Tukey’s HSD posthoc test. Paired two-sample t-tests were used to compare the means in conidia density. The distributions of model residuals were assessed with a Shapiro-Wilk test and QQplots using the “qmath” function of the lattice package in R. As per manufacturer instructions for our haemocytometer, we calculated the average number of conidia per square x the dilution factor (= 10) x 10,000 to acquire conidia cells/ml (each haemocytometer square holds 10**^-^**^4^ ml of the suspension).

We applied raster analyses in Python to assess the growth of conidia while in the Petri dishes. Images were acquired using a Fujifilm XT-4 camera. Images were saved in PNG format and cropped to remove any irrelevant background. Image colour representation was converted from RGB to HSV using the OpenCV library in Python. This conversion was chosen for its ability to separate colour components, providing a more intuitive representation, and greenness was isolated due to the colour of *T. harzianum* conidia. The green colour range in the HSV colour space was defined as [35, 35, 35] to [180, 255, 255]. This range was determined through a combination of literature review and empirical analysis of image characteristics. A binary mask was created by thresholding the images using the defined green colour range. This step resulted in the isolation of regions corresponding to green colour.

The quantification of the green colour involved automated counting of the number of green pixels in the binary mask. The percentage of greenness was calculated (i.e., automated) by dividing the count of green pixels by the total number of pixels in the image. Statistical analyses, including mean and standard deviation estimations, were performed on the quantified green colour data to assess variations across samples. We used the Mann-Whitney U test (Wilcoxon rank-sum test) in R to compare the percentage of green coverage between treatment groups. Data visualisations were produced using a combination of R, Python and Adobe Illustrator Creative Cloud 2022 (Adobe 2021).

## Results

### Weight of teabags before dehydration

Acoustic stimulation at both 70 and 90 dB had a strong effect on increasing *untreated* and heat-treated green teabag biomass compared to controls (untreated: F(3, 58) = 1234.59, *p* = < .001; Eta^2^ = 0.98, 95% CI [0.98, 1.00]; Tukey HSD*, p =* < 0.05; Figure 3a; heat-treated: F(3, 58) = 139.80, *p* = < .001; Eta^2^ = 0.88, 95% CI [0.83, 1.00]; Figure 3a). Acoustic stimulation also had a strong effect increasing untreated and heat-treated rooibos teabag biomass compared to controls (untreated: F(3, 58) = 238.62, *p =* < 0.001; Eta^2^ = 0.93, 95% CI [0.90, 1.00], Tukey HSD, *p =* < 0.05 (Figure 3c); heat treated: F(3, 58) = 179.15, *p* = < 0.001; Eta^2^ = 0.90, 95% CI [0.86, 1.00], Tukey HSD, *p =* < 0.05; Figure 3b).

**Figure 3.**
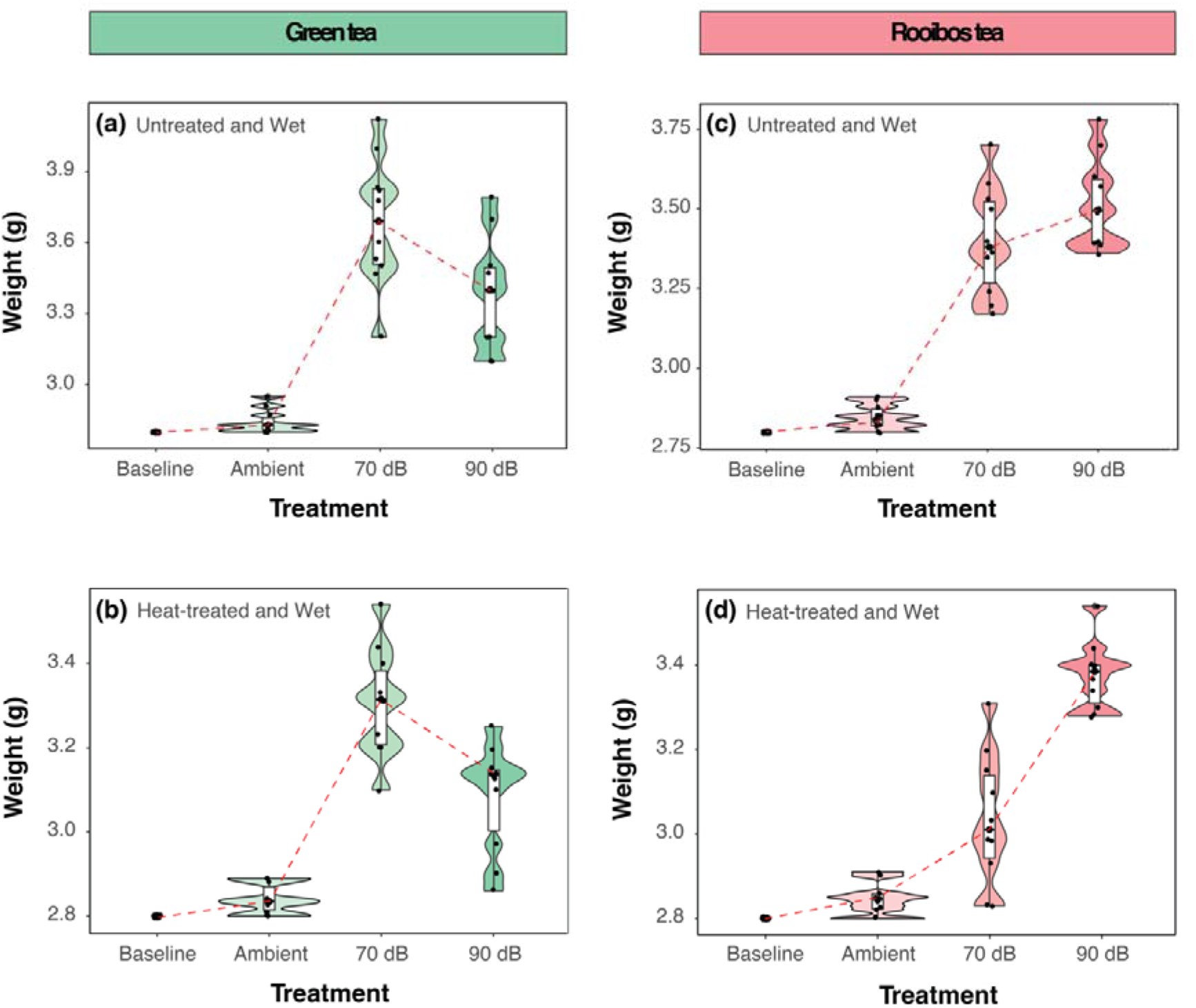
Boxplots of green and red tea weight separated by treatment groups (Ambient control (*n* = 10), 70 dB (*n* = 10) and 90 dB (*n* = 10). Boxplots show values *before* dehydration (i.e., “wet”) for (a) green tea untreated, (b) green tea heat-treated, (c) rooibos tea untreated, and (d) rooibos tea heat-treated. Baseline values (*n* = 30) are shown at the first point of the x-axis (standardised to 2.8 g). Violins (the undulating outline around the boxplots) represent kernel density estimations. Each plot has a red dashed guideline, showing mean trends—these are for visual aid purposes only and were added to the plots using Adobe Illustrator (Adobe Illustrator CC 2023 27.3).

### Weight of teabags after dehydration

Acoustic stimulation at both 70 and 90 dB had a strong effect on increasing *untreated* and heat-treated green teabag biomass compared to controls (untreated: F(3, 58) = 293.01, *p* = < .001; Eta^2^ = 0.94, 95% CI [0.91, 1.00], Tukey HSD, *p =* < 0.05; Figure 4a; heat-treated: F(3, 58) = 1093.40, *p* = < 0.001; Eta^2^ = 0.98, 95% CI [0.98, 1.00], Tukey HSD, *p =* < 0.05; Figure 4b). There was no difference between the 70 dB and 90 dB groups (Tukey HSD, *p* = 0.73).

**Figure 4.**
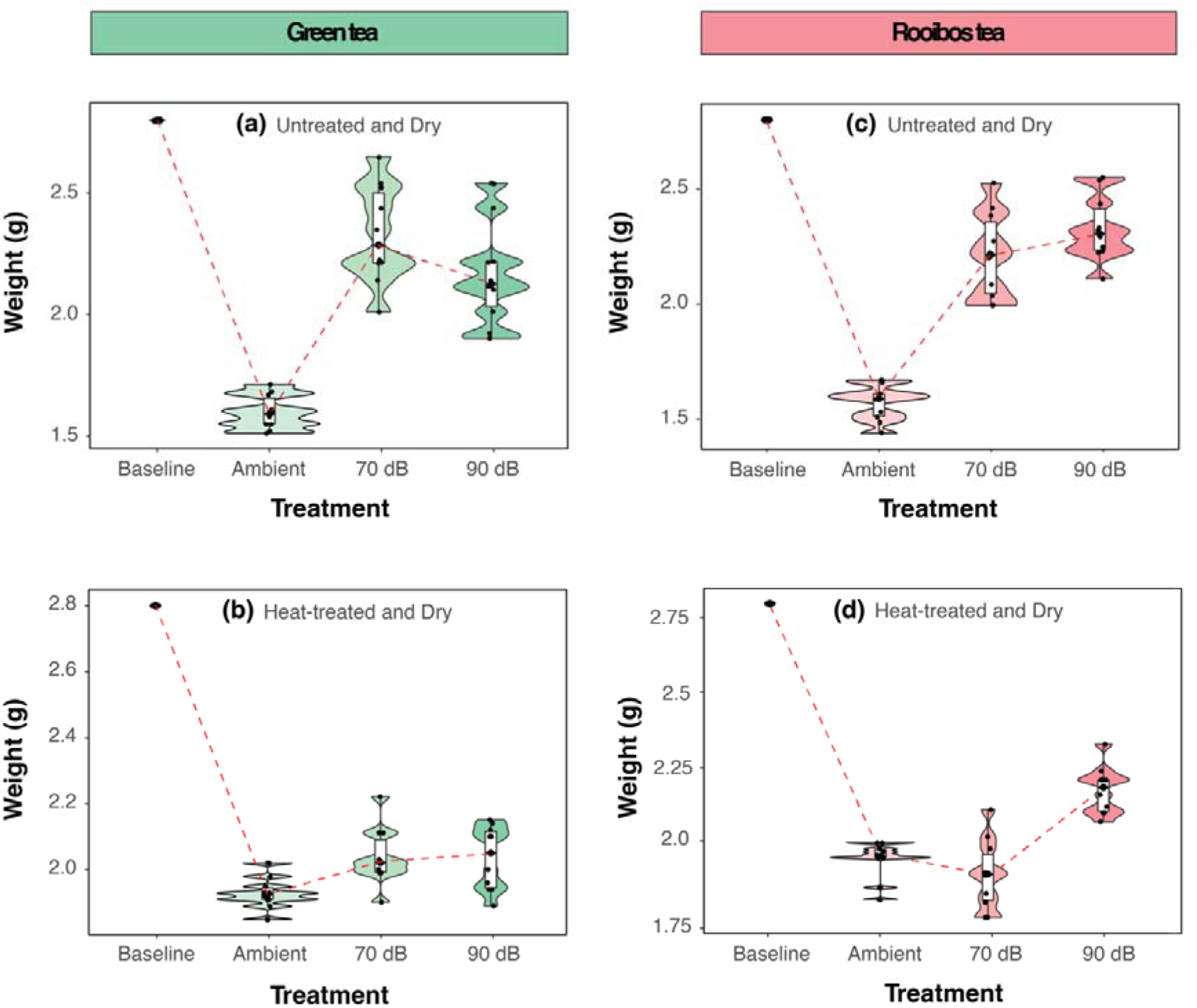
Boxplots of green and red tea weight separated based on treatment groups (Ambient control (*n* = 10), 70 dB (*n* = 10) and 90 dB (*n* = 10). Boxplots show values *after* dehydration (i.e., “dry”) for (a) green tea untreated, (b) green tea heat-treated, (c) rooibos tea untreated, and (d) rooibos tea heat-treated. Baseline values (*n* = 30) are shown at the first point of the x-axis (standardised 2.8 g). Violins (the undulating outline around the boxplots) represent kernel density estimations. Each plot has a red dashed guideline showing mean trends—these are for visual aid purposes only.

Acoustic stimulation at both 70 and 90 dB had a strong effect on increasing *untreated* rooibos teabag biomass compared to controls (untreated: F(3, 58) = 432.87, *p* = < 0.001; Eta^2^ = 0.96, 95% CI [0.94, 1.00]), Tukey HSD, *p =* < 0.05; Figure 4c). There was no difference between the 70 dB and 90 dB groups (Tukey HSD, *p* = 0.34). Acoustic stimulation at 90 dB had a strong effect on increasing *heat-treated* rooibos teabag biomass compared to controls (heat-treated: F(3, 58) = 915.07, *p* = < 0.001; Eta^2^ = 0.98, 95% CI [0.97, 1.00]), Tukey HSD, *p =* < 0.05; Figure 4d). There was no difference between the 70 dB and the ambient (control) group (Tukey HSD, *p* = 0.11).

### Visual assessment of fungal biomass

Fungal biomass was not visibly present in any teabags at the start of the experiment. After 14 days of acoustic stimulation, fungal biomass was visibly abundant in the 70 dB and 90 dB treatment groups, for both green tea and rooibos teabags, and on both the interior and exterior of each teabag (Figure 5). Fungal biomass was less visible in the 20 control teabags. The internal biomass of the teabags in the 70 dB and 90 dB treatment groups was dense compared to the control (which had clear space between the leaves and bag).

**Figure 5.**
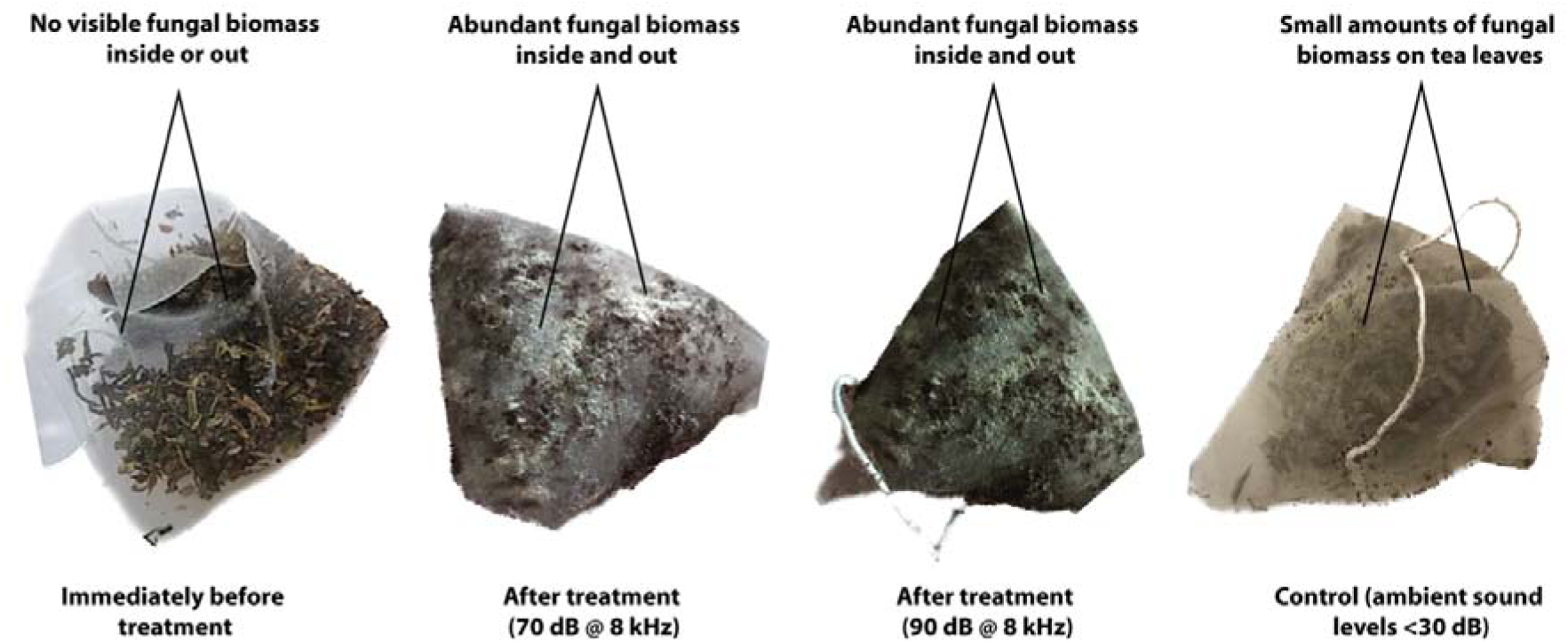
Fungal biomass was visibly absent from the teabags before the treatment of acoustic stimulation. However, teabag mass increased considerably under 70 dB and 90 dB treatments (and no inter-treatment differences), particularly in the non-heat-treated group (pictured), with fungal biomass visibly abundant inside and outside of the teabag netting. The density of mass within the teabags is also visible when compared with the ‘before treatment’ teabags and the control. The control sample showed small amounts of fungal growth; however, this was limited to tea leaves. These visual signs were consistent across untreated and heat-treated samples.

### Soil pH

There were no significant changes in soil pH between the beginning and the end of the experiment for any treatment group. However, dehydration had a weak effect on increasing soil pH (heat-treated soil pH xL = 6.90, SD = 0.04, untreated soil pH xL = 6.94, SD = 0.04, t = -5.03, df = 29, *p* = <0.05).

### Radial (mycelial) growth

Acoustic stimulation had a strong effect on increasing mycelial radial growth at day two (acoustic treatment: xL = 60.5 mm, SD = 3.09; control: xL = 58.5 mm, SD = 1.89; t = 2.5, df = 18, *p* = 0.02). On day three, there was no effect of acoustic stimulation on mycelial radial growth (t = 0.5, df = 18, *p* = 0.58). However, by day four, there was a strong effect of acoustic stimulation and mycelial growth had increased substantially (acoustic treatment: xL = 89.5 mm, SD = 1.07; control: xL = 82.8 mm, SD = 8.5; t = 3.66, df = 18, *p* = 0.001). By day five, there was again a strong effect of acoustic stimulation on mycelial radial growth (acoustic treatment: xL = 89.6 mm, SD = 1.07; control: xL = 83.4 mm, SD = 7.8; t = 3.37, df = 18, *p* = 0.003).

### Conidia growth (proxy)

Acoustic stimulation had a strong effect on increasing conidial growth (Figure 6; day five acoustic treatment: xL = 2.8% coverage, SD = 2.9; control: xL = 0.39% coverage, SD = 0.94; W = 61.5, df = 18, *p* = 0.001).

**Figure 6.**
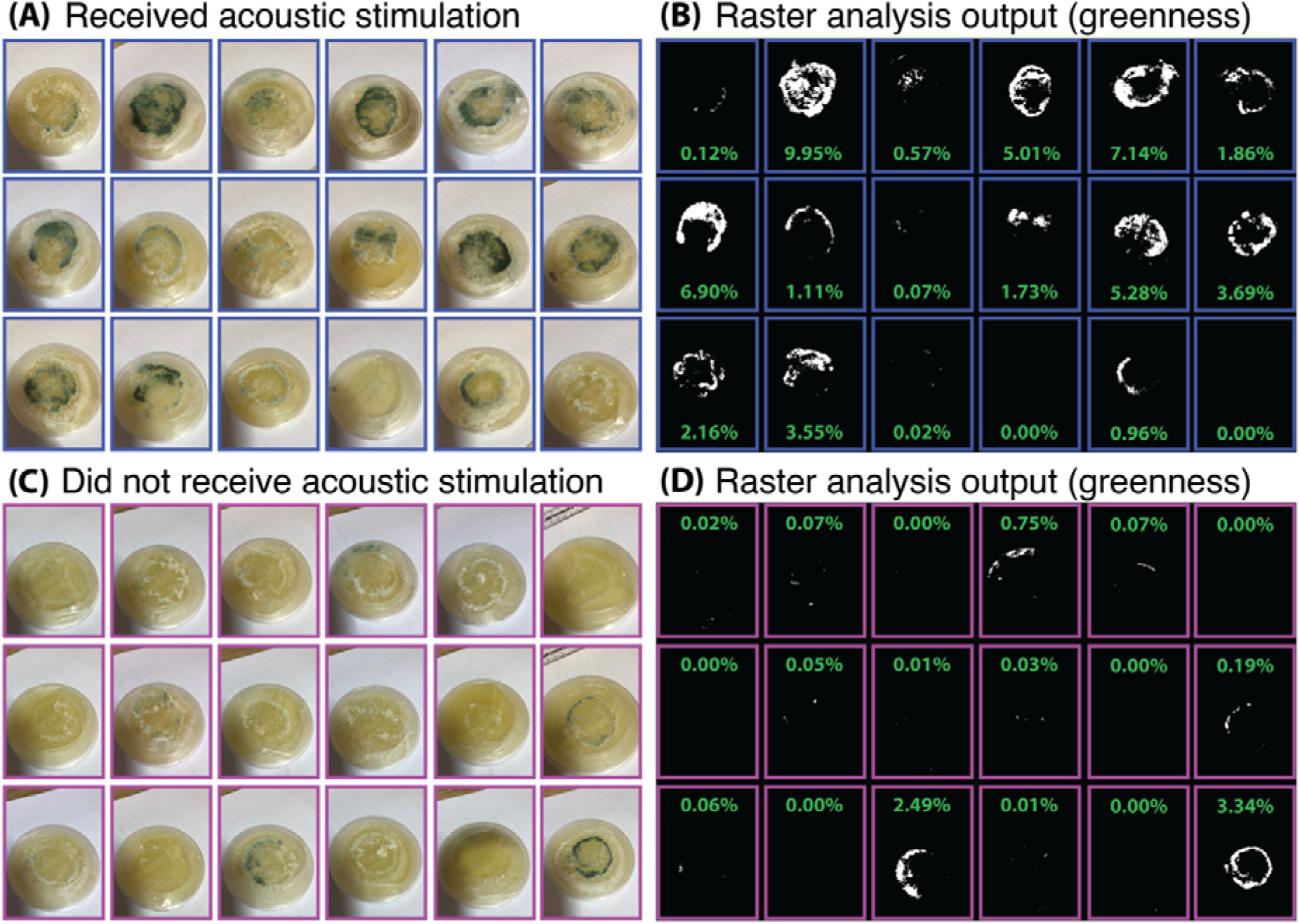
Images of the Petri dishes containing *T. harzianum* culture on day 5 and the outputs of the raster analysis of greenness (A = acoustic stimulation group, B = acoustic stimulation group, C = culture control group, D = raster analysis output for the control group) – including the percentage of green cover as a proxy for conidia growth.

### Conidia cell density

Acoustic stimulation had a strong effect on increasing conidial density (day five acoustic stimulation: conidial density: xL = 2,421,052 cells/ml; control: xL = 542,105 cells/ml; t = 18.2, df = 18, *p* = <0.001). Cell density was strongly and positively correlated with the percentage of green cover in the Petri dishes (r_s_ = 0.63, S = 2855, *p* = <0.001; Figure 7).

**Figure 7.**
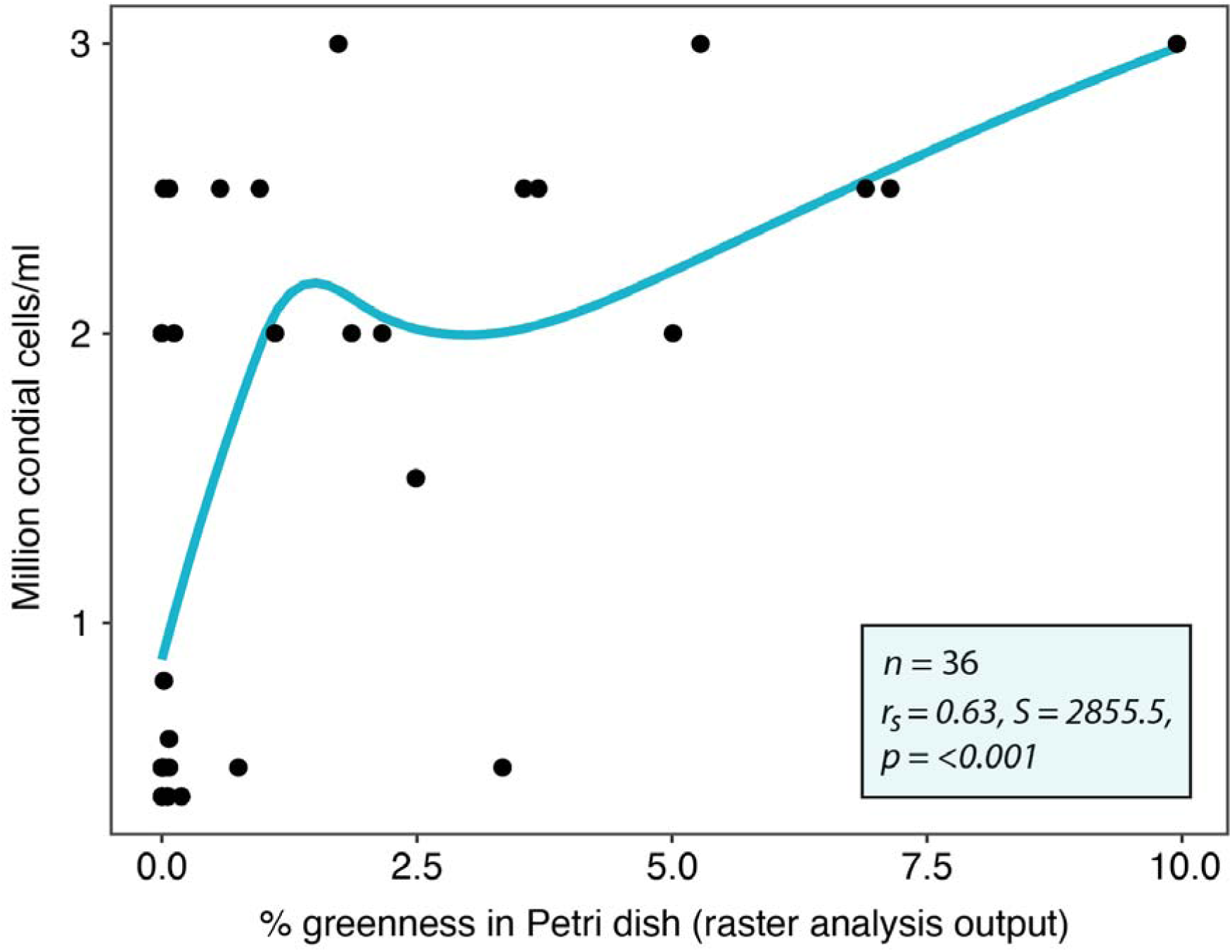
Correlation between the conidia cell density (as determined via the haemocytometer) and the percentage greenness coverage in the Petri dishes. The blue line represents a smoothing (direction and strength of correlation) fitted to the data points.

## Discussion

Sound is a critical component of ecosystems, and we can detect acoustic properties to monitor the restoration of soil biodiversity (Robinson et al. 2023). However, the application of acoustic properties in a targeted way to alter and potentially enhance soil restoration processes remains unexplored. Our study showed that acoustic stimulation increases fungal biomass and aspects of decomposition in an experimental soil mesocosm setting, and enhances the activity of a plant growth-promoting fungus in a laboratory setting. These preliminary results serve as a foundation for extending research into sonic restoration (e.g., exploring the effects of specific acoustic parameters on particular fungal species and/or communities), plus the mechanisms by which soil life is affected by sound (e.g., piezoelectric effects to and/or mechanoreceptor stimulation of cellular and/or molecular processes). There is potential to use this technology to improve ecosystem restoration outcomes, as well as agricultural and clinical settings.

### Acoustic stimulation increases soil fungal biomass

We show in mesocosm experiments that our acoustic treatments increased the mass of green and rooibos teabags. Our sound parameters (70 dB and 90 dB @ 8 kHz) altered fungal biomass most likely by increasing their organic matter content due to stimulating fungal growth and/or moisture absorption. We suggest that the fungi within acoustic treatments were decomposing organic matter (i.e., the tea) and gaining weight faster than controls – i.e., they held more water than energy lost as heat, compared to controls.

Piezoelectric effects, induced by mechanical pressure (e.g., from acoustic waves) on piezoelectric materials, can influence cellular and molecular processes in living organisms, including microbiota (Gazvoda et al. 2022). Mechanoreceptor stimulation, such as the activation of mechanosensitive ion channels in cells (e.g., by touch, sound and other mechanical stimulation), plays a pivotal role in translating mechanical signals into cellular responses, impacting processes like gene expression and cell signalling pathways (Sun et al. 2022). Acoustic stimulation can also affect the production of various metabolites in *Saccharomyces cerevisiae* yeast in a liquid medium (Shah et al. 2016; Harris et al. 2021). It can also influence the production of quorum sensing-regulated pigments, prodigiosin and violacein (Shah et al. 2016). Therefore, with refinement, acoustic stimulation has the promise to be developed into a tool to affect specific ecological functions (e.g., organic matter decomposition). Our results are consistent with previous studies, including Hofstetter et al. (2020), who showed that refrigerator acoustic vibrations can increase fungal biomass, and Harris et al. (2021), who found that 90 dB acoustic stimulation increased fungal growth in liquid media. Increased fungal biomass in our acoustic stimulation treatments was also supported by the visual inspection of our experimental tea bags.

We do note some inconsistent findings. The heat-treated 70 dB rooibos group was lighter than the baseline but heavier than the ambient control group after dehydration. The cause of this reduced biomass is unknown, but was potentially due to this type of acoustic stimulation increasing organic matter decomposition in the woodier rooibos tea when microbial communities have been degraded (e.g., by our heat-treatment), compared to 90 dB and the leafier green tea.

### Acoustic stimulation increases the activity of plant growth-promoting fungi

We show that acoustic stimulation increased the growth rate and sporulation of *T. harzianum*, a well-known plant growth-promoting fungus (Lòpez et al. 2023). Our novel raster analysis provided a good measure of conidia growth/coverage in Petri dishes and the haemocytometer. The potential mechanisms causing such effects may also be piezoelectric and mechanoreceptor stimulation, but this needs further investigation. Our results are consistent with Hoffstetter et al. (2020), who showed fungal growth increases at high frequencies (above 5 kHz, as per our study). This study also suggested that low frequencies (below 165 Hz) could reduce the growth rate of *Botrytis sp*.

Whether certain sound parameters can target particular fungal species or guilds is yet to be determined. This is a worthwhile research enquiry because it could have broad-reaching implications, such as improving ecosystem restoration and agricultural outcomes (e.g., increasing the biomass of desirable fungi including plant growth-promoting and commercial species, suppressing undesirable fungi such as pathogens humans and desirable plants). Of course, the potential unintended or undesirable consequences of using this technology need to be investigated (e.g., non-target impacts).

In an ecosystem restoration context, we suggest two priority applications to further develop: (1) applying acoustic stimulation to enhance the production efficiency of microbial inoculants (e.g., potentially enhancing the growth rate but also the viability, quality and functional potential of beneficial fungal spores), and (2) the direct application of a sound source in ecosystems (*in-situ*) to help improve their biological integrity via a direct effect on soil and potentially non-soil microbiota. While still in the early stages, our results are encouraging to develop innovative restoration techniques that leverage sound to alter soil ecosystem functioning. Considering the broader restoration imperative, exploring the role of acoustic stimulation represents an exciting and underexplored avenue of research. Expanding our understanding of the relationships between acoustics, soil microbiota, and ecosystem functioning paves the way for advancements in restoration and microbial ecology.

## Conclusion

Our study introduces a novel dimension to the soil restoration domain by investigating the effects of acoustic stimulation on fungal biomass and plant growth-promoting fungi. Demonstrating a tangible impact on fungal activity, our findings suggest that carefully tuned acoustic parameters can influence soil (and potentially plant) components via their effect on fungi. We propose two critical avenues for future research: optimising acoustic stimulation for microbial inoculants for plants and exploring in-situ applications to enhance biological integrity and desirable processes in eco- and agro-systems. Despite the need for further investigation into potential unintended consequences, our study marks an important stride toward leveraging sound as a tool for innovative and effective soil ecosystem restoration.

## Acknowledgements

MFB was funded by the Australian Research Council [grants LP190100051, LP190100484, DP210101932] and the New Zealand Ministry of Business Innovation and Employment [grant UOWX2101].

---

**Table S1.** Treatment groups in this study (Aim 1) applied to 10 pots of heat-treated and untreated soils.

| Amplitude | Frequency | Daily duration | Number of days |
| --- | --- | --- | --- |
| 70 dB | 8 kHz | 8 hrs | 14-days |
| 90 dB | 8 kHz | 8 hrs | 14-days |
| Ambient (<30 dB) | Ambient | 8 hrs | 14-days |

